# Transformer-Based Phenotyping of Rice Root Aerenchyma Across Environments Enables Climate-Smart Rice Selection

**DOI:** 10.64898/2026.01.30.702889

**Authors:** Hani Atef, Laura Fierro-Dominguez, Paula Andrea Lozano-Montaña, Sergi Navarro Sanz, Julie Bals, Benoît Clerget, Christophe Périn, Maria Camila Rebolledo, Romain Fernandez

## Abstract

Quantification of root anatomical traits such as cortical aerenchyma is key to understanding rice adaptation to diverse water regimes. Recently, the role of aerenchyma in regulating methane emissions has been demonstrated, making it a target for climate change mitigation. Despite its importance, breeding for root anatomical traits remains limited because manual analysis of root cross-sections is labor-intensive, inconsistent, and poorly scalable, and analysis pipelines do not generalize across heterogeneous imaging conditions. We present a deep learning pipeline based on a recent vision transformer architecture to automatically segment rice root anatomical structures and quantify aerenchyma. The model was trained on a multi-environment dataset of 1,760 annotated rice root cross-sections acquired across growth stages, cultivation systems, and countries, using a collaboratively defined annotation protocol. The model achieved high segmentation performance (mean Intersection-over-Union > 0.92) and near-perfect aerenchyma ratio quantification (R^2^ = 0.98), and was evaluated by two experts as performing on par with, and in some cases better than, expert annotators.

Delivered as open-source software with an online interactive demonstrator, the pipeline revealed differences in aerenchyma across genotypes, water regimes, environments, and developmental stages. Overall, this work demonstrates that transformer-based segmentation enables high-throughput anatomical phenotyping, supporting scalable and climate-smart rice breeding.

**HIGHLIGHTS:**

- Transformer-based segmentation enables robust aerenchyma phenotyping across environments
- A SegFormer model achieves expert-level accuracy on diverse rice root cross-sections
- Automated analysis delivers near-perfect lacuna-to-cortex ratio quantification (R^2^ ≈ 0.98)
- Our online demonstrator supports scalable, climate-smart rice breeding applications

## INTRODUCTION

Cortical root traits play a central role in climate adaptation and resilience in Poaceae (Jones et al., 2025a), highlighting the need for phenotyping tools that enable their effective incorporation into breeding programs. Besides, quantifying anatomical traits in roots is instrumental for modelling plant water uptake, gas transport, and adaptation to environmental constraints (Atkinson et al., 2019; Lynch, 2019). In roots, aerenchyma is a cortical tissue characterized by enlarged gas spaces exceeding typical intercellular voids, playing a dual role in rice (Suralta and Yamauchi, 2008). Its presence supports oxygen and methane transport under flooded conditions (Jiménez and Pedersen, 2023) or can reduce root metabolic cost under drought (Caetano et al., 2025). But its presence can also affect the radial water conductivity under non-flooded aerobic conditions (Song et al., 2023). As a result, accurate and reproducible measurement of aerenchyma, cortex area, and the central cylinder could become essential for studies aiming to identify genotypic variation in anatomical plasticity and to support breeding programs targeting climate adaptation and resilience.

Several tools have been developed to extract anatomical traits from root cross-sections. Early semi-automatic pipelines such as RootScan (Burton et al., 2012) and PHIV-RootCell (Lartaud et al., 2015) rely on rule-based image processing implemented in ImageJ, requiring user-guided segmentation steps and threshold tuning. These methods enable the quantification of features such as stele diameter, cortex width, and vessel number, and can indirectly inform estimates of aerenchyma proportion. However, they are sensitive to image variability and remain dependent on manual supervision, which limits their throughput and reproducibility, especially across sampling variability, including the developmental stage of the roots, the species, and the culture conditions. RootScan was developed and validated primarily using crown roots from young plants of maize and, to a lesser extent, rice and bean (25-28 days after sowing) grown under relatively controlled and field conditions. By contrast, PHIV-Rootcell was designed for analysing very young rice seedlings (6 days after sowing) that are grown under highly controlled conditions on a Murashige and Skoog medium, focusing on the root tip cross-section of developing radicles. RootAnalyzer (Chopin et al., 2015) further extends automated anatomical analysis to a later developmental stage, having been validated on primary roots of wheat (Triticum aestivum L.) genotypes grown under contrasting water regimes (well-watered and drought), with cross-sections obtained approximately 4 cm from the root tip at 50–56 days after sowing. While these earlier tools advanced the field, segmentation of mature cortical aerenchyma across diverse environments poses distinct challenges that require complementary approaches, targeting data source diversity and robustness to varying observation conditions.

In recent years, numerous deep plant phenomics tools have emerged (Ubbens and Stavness, 2017), and root anatomics make no exception. RootAnalyzer (Chopin et al., 2015), which applies contour-based segmentation with fixed pipelines, and DL-RootAnatomy (Wang et al., 2020), which uses object detection networks (Faster R-CNN) to identify anatomical regions, are representative examples. However, while these approaches reduce manual intervention, they are trained on controlled laboratory images, so their robustness under more variable data from field growing setups remains uncertain, despite being required for leveraging phenotyping for studying target systems. Furthermore, none of the existing tools was specifically designed to segment the highly variable cortical lacunae. For instance, RootAnalyzer only provides a rudimentary classification of large cortical air spaces, and DL-RootAnatomy does not address them explicitly. Ultimately, these tools have not yet been applied to distinguish anatomical differences across genotypes, treatments, or developmental stages in practice, limiting their possible applicability to support ongoing breeding efforts. However, recent efforts in acquisition systems have pushed anatomical phenotyping possibilities toward higher throughput using custom-built platforms such as the Rapid Anatomics Tool (RAT) (Jones et al., 2025a), including an efficient and scalable mechanical sectioning design, combined with a semi-automatic analysis pipeline. And high-resolution 3D imaging approaches, such as Laser Ablation Tomography (LAT), provide access to volumetric anatomy (Lynch et al., 2021) but involve costly instrumentation located at specific places, limiting its practical deployment across large, field-derived populations, and requiring custom 3D analysis solutions.

Together, these tools highlight a gap in AI-driven plant research: the lack of automated methods fitting with the standard laboratory imaging and computing environment, capable of quantifying aerenchyma formation in mature zones of seminal roots, where cortical air spaces are highly heterogeneous and influenced by genotype, environment, and developmental stage. In this work, we present a robust and scalable pipeline for anatomical segmentation of root cross-sections using transformer-based deep learning methods. Our method focuses specifically on the cortex and cortical aerenchyma. It is trained on a multi-environment, field-derived dataset of nearly 1800 curated images and requires no parameter tuning. Built on the SegFormer architecture (Xie et al., 2021), our model accurately segments the cortex and lacunae, achieving expert-level performance across heterogeneous acquisition conditions, which we assessed through an independent expert review, confirming that model errors are fewer and less biologically impactful than those in manual annotations. The pipeline is fully open-source, includes a web-based interface for non-specialist users, and is routinely used in ongoing phenotyping campaigns. Finally, we demonstrated how this tool is able to differentiate rice cortical aerenchyma across different water treatments, genotypes, growth stages, and growth environments (from hydroponics to field). By providing a fast, reproducible, and field-validated method for quantifying aerenchyma, this work contributes a practical tool for large-scale anatomical phenotyping and supports breeding efforts targeting climate-smart rice systems.

## MATERIALS AND METHODS

### 2.1 Plant material and imaging

Root cross-sections were obtained from multiple experiments conducted in different countries (Colombia and France) and environments (hydroponics, petri dishes, pots, and field), see Fig. 1 and Table 1. Root segments were collected from either seminal roots (hydroponic) or crown roots (soil or pot) and excised at either 1, 2, or 3 centimeters from the root tip (hydroponic) or 1 cm from the root base (soil or pot).The global dataset (across use-cases) corresponds to different root maturity levels (and lengths), thus the dataset includes a representative variability of section position relatively to the tip, and hence, a continuum of aerenchyma maturation levels.

**Table 1.** Summary of the use-cases providing images and annotations, including characteristics of their experiments.

| Use-case code | Country | Rice varieties | Growth stage | Growing conditions | Research group | Experimenter |
| --- | --- | --- | --- | --- | --- | --- |
| UC1<br>-JUL | France | Mutants from a background of Kitaake | 5 leaves | Hydroponic in a growth chamber | CIRAD/DARS | Julie Bals |
| UC2<br>-HYDRO | Colombia | Panel of 20 Indica, Japonica, Aus, O. rufipogon, and Tropical japonica | 6-7 leaves | Hydroponic in a glass greenhouse | CIAT | Paula Lozano |
| UC3<br>-SHELTER | Colombia | Panel of 19 Indica, Japonica, Aus, O. rufipogon, and Tropical japonica | 6-7 leaves and panicle initiation | Rain-out shelter with soil, flooded / not flooded | CIAT | Paula Lozano |
| UC4<br>-MUT | France | Mutants from the Kitaake genotypes | 5 leaves<br>-1cm base | Growth chamber, pots with substrate | CIRAD/<br>PHENOMEN | Domitille Tur |
| UC5<br>-VARIETIES | France | Panel of 10 Indica and japonica tropical | 5 leaves | Greenhouse, pots with soil, flooded / not flooded | CIRAD/<br>PHENOMEN | Justin Savajols |
| UC6<br>-LFD | France | IR64 (indica) and Primavera (japonica) | 5 leaves | Greenhouse, pots with soil, flooded / not flooded | CIRAD/<br>PHENOMEN | Laura Fierro Dominguez |

**Fig. 1.**
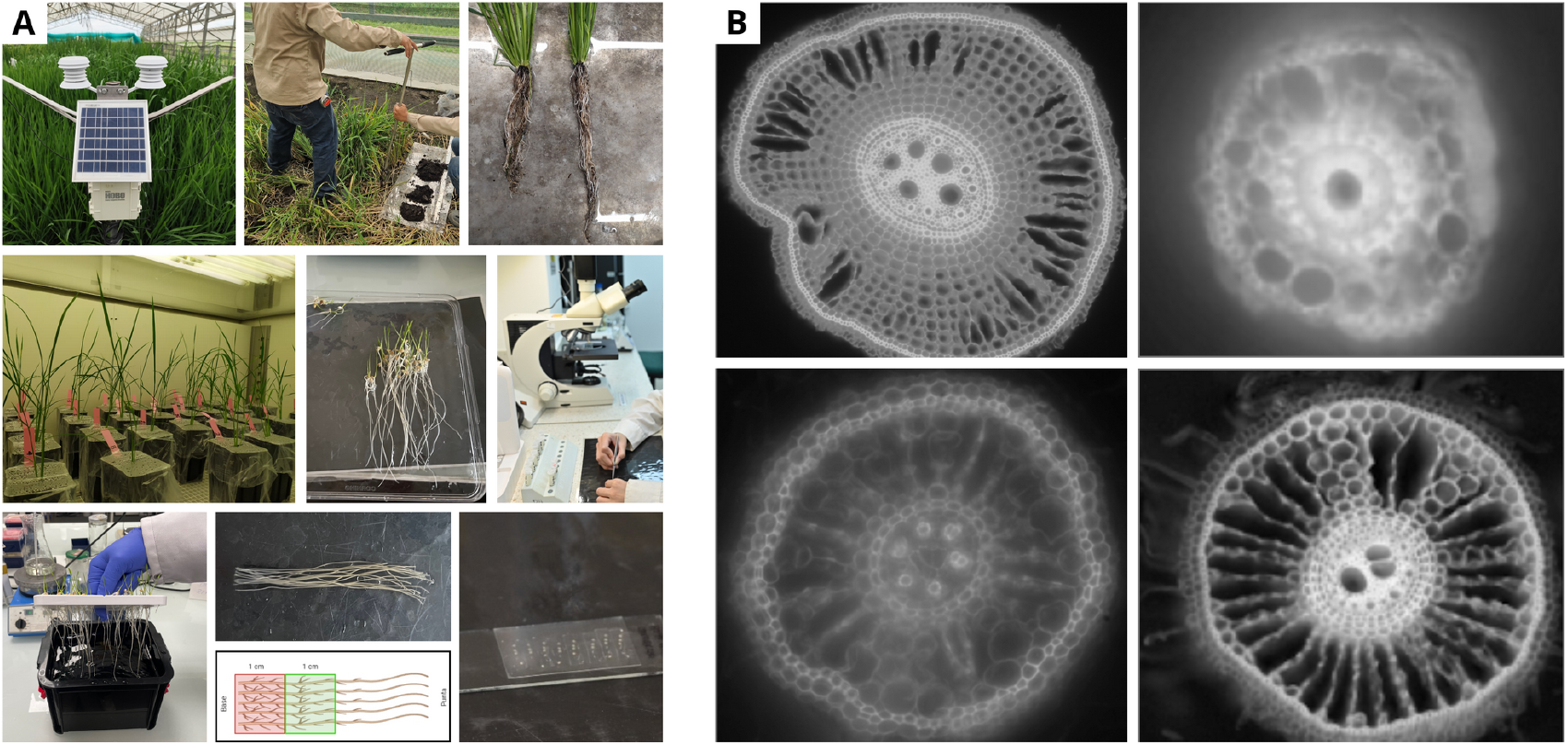
Culture environments, imaging setups, and resulting root cross-section images. (A) Overview of culture environments and imaging systems. (B) Representative images obtained from the experimental contexts. **ALT-TEXT:** Figure showing the diversity of experimental contexts used in the study. Panel A illustrates different rice growth environments and corresponding imaging setups. Panel B displays representative rice root cross-section images obtained under these conditions, highlighting variability in image appearance across environments.

The segments were embedded in 5% agar and sectioned transversely using a vibratome, producing uniform histological slices for anatomical analysis. Imaging was performed using epifluorescence microscopy under controlled acquisition settings. Illumination intensity was maintained at 85%, and images were captured at magnifications ranging from 10x to 20x, depending on root diameter. All images were acquired as 8-bit grayscale TIFF files with sufficient resolution to visualize cortical tissues, the endodermis, cell walls, and lacunae. The resulting dataset comprised 2,498 raw microscope images distributed across several acquisition batches with heterogeneous quality (Fig. 1). Detailed metadata (site, genotype, treatment, acquisition date, magnification) were kept to assess robustness across varying conditions.

### 2.2 Image quality control and dataset curation

To exclude damaged or low-quality images, we developed a custom Python graphical interface enabling rapid visual inspection and triage of all samples. Images were manually flagged for removal based on the following criteria: blur or defocus artifacts on a large portion of the slice (Fig. 2C), incomplete or truncated root sections or additional structures (Fig. 2A, 2C, 2D, 2E), topological artifacts obstructing the cortex (Fig. 2D), and extreme shape deformation or anatomical ambiguity preventing reliable annotation (Fig. 2D, 2F). These categories cover many sources of image defects, including the presence of large water droplets preventing visualization of part of the slice; one or multiple lateral roots inducing the presence of “support tissue” altering the local aerenchyma distribution; and large concavities in the slices, indicative of damaged roots and unreliable measurements of cortex surface. After the manual flagging step, the tool generated a full audit report (CSV), which was used to give feedback to the image providers to enable discussion and improvement. After this curation step, 1,760 images (70.46% of the original dataset) remained for model development.

**Fig. 2.**
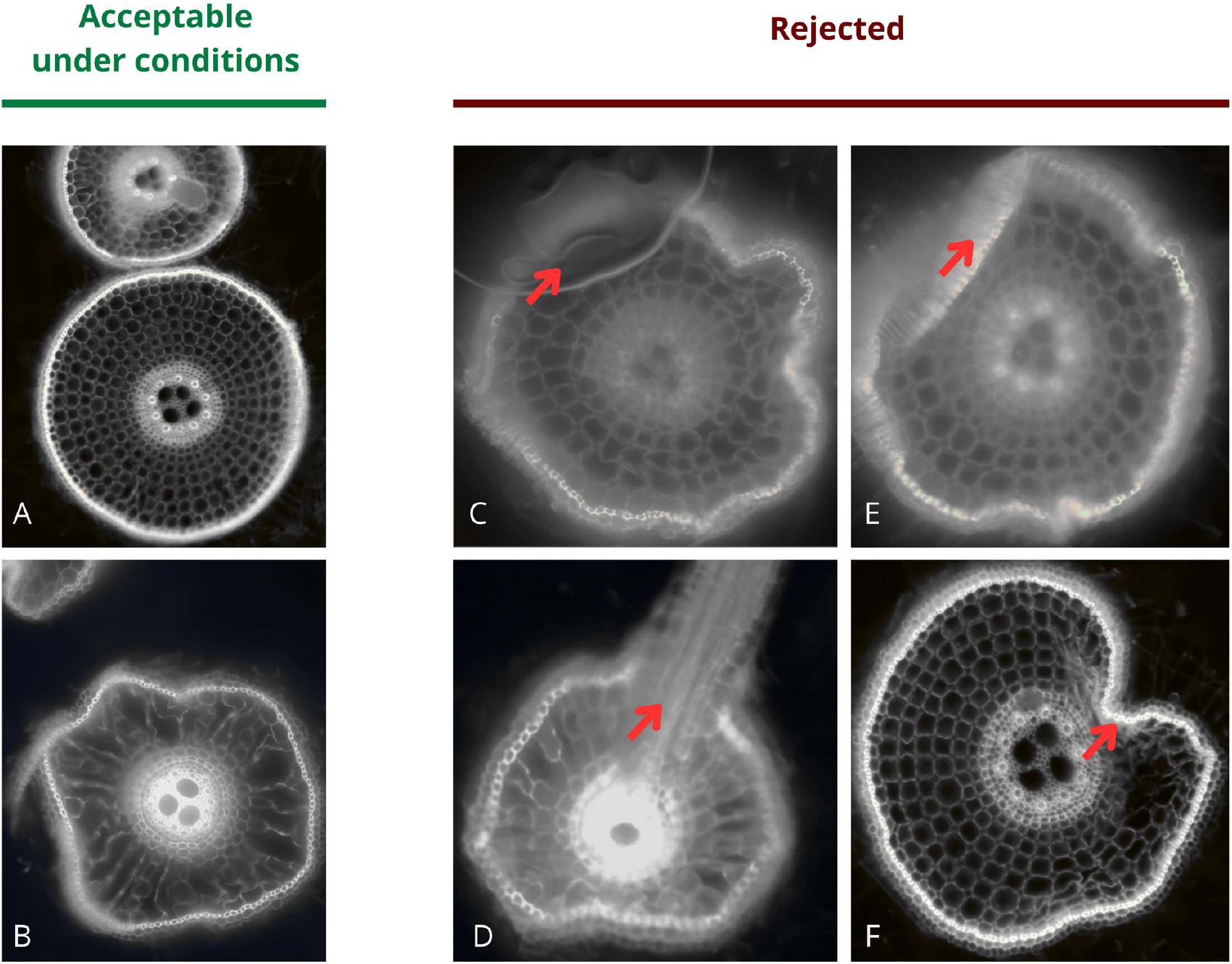
Representative examples of curated rice root cross-section images. A: a correct root cross-section, surrounded by an additional object, that will be automatically removed during processing after cortex segmentation by selecting the bigger cortex region. B: limit of circularity for accepting a cross-section, to foster an acceptable level of biological meaning in the measurements. C, D, E, F, cross-sections presenting at least one important defect, thus being rejected from the set during manual screening (C: water drop, D: presence of a lateral root, E: the slice is cut, F: the slice is damaged, probably by manipulation). **ALT-TEXT:** Figure illustrating the curation criteria applied to rice root cross-section images. Panel A shows a valid cross-section with an adjacent object that is removed automatically during processing. Panel B illustrates the circularity threshold used to retain biologically meaningful sections. Panels C–F show examples of cross-sections rejected during manual screening due to major defects, including the presence of a water drop, a cut section, a lateral root emerging, and a large intrusion of sclerenchyme into the cortex, likely due to a manipulation error.

### 2.3 Target description

#### Aerenchyma genesis and visual phenotype

Our image analysis objective is the identification and quantification of cortical aerenchyma, the air-filled cavities that develop within the cortex. The aerenchyma observed here is lysigenous, arising from programmed cell death and the progressive lysis and collapse of specific cortical cells, which ultimately generate the interconnected air spaces that facilitate internal aeration under hypoxic stress. Aerenchyma development is gradual, and our dataset captures a continuum of maturation stages. In our epifluorescence cross-sections, aerenchyma is not revealed by a specific stain; instead, images mainly show general cell-wall autofluorescence, so both living cells and air spaces appear as “holes” bounded by fluorescent walls. Consequently, aerenchyma identification relies on a combination of morphological cues rather than a dedicated intensity signature. In early stages, incipient aerenchyma can be subtle: healthy cortical cells are typically polygonal and tightly packed, whereas the first lysing cells exhibit tortuous or wavy walls, local wall discontinuities, and partially emptied lumens (Fig. 3A-3B). With progression, adjacent lacunae expand and coalesce into larger cavities. At advanced stages, these cavities are often larger than individual cells and become elongated along the root radius (centre-to-periphery), consistent with the fact that mature aerenchyma can span what would normally correspond to multiple radial cortical cell layers. Fig. 1B illustrates representative early and late aerenchyma morphologies, ranging from small irregular lacunae to large continuous air channels traversing the cortex.

**Fig. 3.**
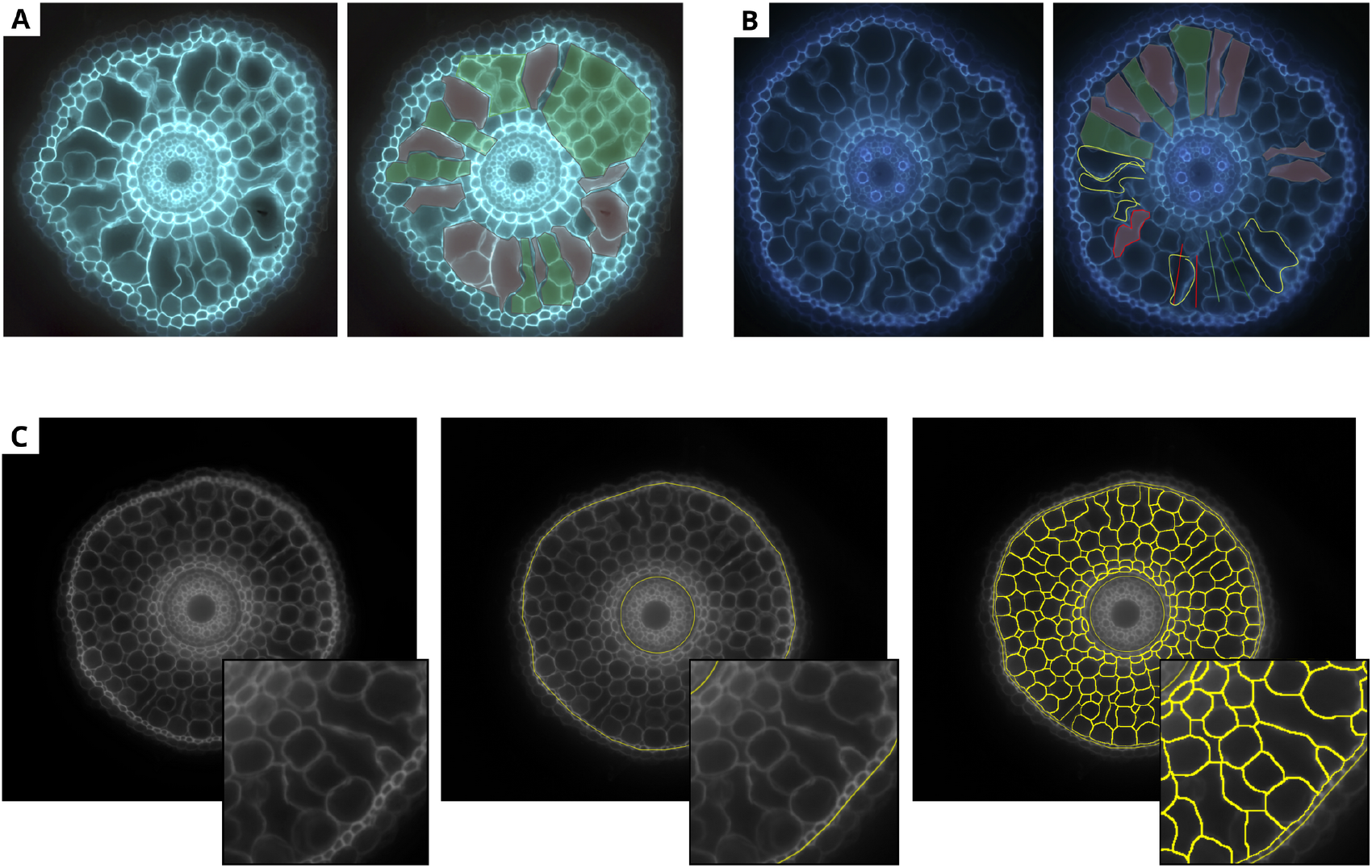
A and B: collaborative target descriptions for challenging images (extracts from the annotation guide). On the left: aerenchyma are indicated in red and non-aerenchyma in green, based on a previous consensus-building meeting. On the right: an ongoing discussion within the annotators team on an online shared document, conducted before the start of the annotation task. In the image, the yellow lines show a proposed aerenchyma annotation by one annotator, while the other interacted by sharing comments. C: steps of the manual annotation workflow, from left to right. Left: original image. Middle: manual identification of inner and outer cortical bounds. Right: manual identification of aerenchyma on the automated watershed segmentation using the “Click Aerenchyma” script. **ALT-TEXT:** Figure describing the collaborative annotation strategy and manual annotation workflow. Panels A and B show excerpts from the annotation guide for challenging cases: aerenchyma and non-aerenchyma regions are distinguished following consensus discussions, and an example of annotator interaction on a shared document is illustrated with lines and contours being drawn live onto a raw image during the discussion. Panel C presents the manual annotation workflow, from the original image to cortex boundary definition and final aerenchyma identification by clicks of the annotator.

#### Morphological variability and annotation challenges

The morphogenetic trajectory creates practical challenges for annotation. Early-stage aerenchyma may be difficult to separate from normal cell lumens. In contrast, late-stage aerenchyma may blur conventional notions of “cell boundaries” because collapsing walls and coalescence can make neighbouring compartments appear merged. To a human observer, advanced aerenchyma may even appear as if an entire radial file of cortical cells is missing, with surrounding cells seemingly “fusing” across the cavity. Capturing such variability consistently in annotations is essential because the segmentation decisions in the intermediate stages of the automated pipeline have a large downstream impact on the extracted quantitative traits (the aerenchyma proportion within the cortex section). To mitigate inter-annotator variability, we developed a detailed annotation guide (selected insights in Fig. 3C). The guide provides explicit criteria for identifying aerenchyma at different stages, and pairs textual rules with visual examples highlighting key cues (wall waviness, partial wall lysis, merged intercellular spaces, and radial elongation). This design was motivated by evidence that annotation errors in biomedical image analysis are often driven less by annotator skill (or lack of) than by ambiguous or underspecified labelling instructions, and that concrete visual examples can substantially improve consistency and accuracy, see (Rädsch *et al*., 2023) for an exhaustive work on that point. By integrating these best practices into our protocol, we aimed to maximize inter-annotator agreement and produce higher-quality ground truth for training and evaluation.

### 2.4 Manual annotation workflow

Ground-truth masks were generated through a semi-automated Fiji/ImageJ annotation workflow with annotators using the target description guide and the workflow tutorial. The workflow was divided into a sequence of manual actions over all images, followed by a sequence of automated actions, in order to streamline annotations across a large number of images. Overall, the manual operations required three minutes per slice, making the full dataset a 90-hour annotation task that was distributed over multiple operators and multiple laboratories.

Starting from the raw image (Fig. 3-C, left), annotators first delineated the inner and outer cortical boundaries (Fig. 3-C, middle), using custom macros based on polygon tracing. A watershed-based cell segmentation step (Fig. 3-C, right) was then applied to highlight internal structures, which helped identify air-filled lacunae and cytoplasm within the cortex. Using a dedicated tool “Click aerenchymes”, annotators manually identified among all the segmented structures those corresponding to aerenchyma, following the annotation guide. A morphological closing operation was then applied to fuse potential aerenchyma subregions resulting from oversegmentation errors. The workflow included image verification steps to help annotators flag images that require re-annotation. After the final step, the workflow produced segmented images comprising three classes: 1. background and central cylinder, 2. cortical lacunae, and 3. cortex-endodermis region. These masks were exported as binary TIFF files and served as ground truth for model training.

### 2.5 Data preparation and augmentation

To facilitate an efficient and deployable pipeline, we downscaled all images from the original definition (∼2000×2000) to a fixed target definition of 512×341 pixels. While this downscaling inevitably sacrifices some fine detail visible to human observers, it has been shown in (Ferreira *et al*., 2025) that moderate image downsampling can maintain or even improve a network’s accuracy by providing more contextual information relative to each pixel, because extremely high resolutions contain detail beyond the effective receptive field of the network (leading to fragmented predictions), whereas a lower resolution can better align with the object size and network context window. In this context, we opted for whole-image downscaling rather than dividing each image into high-resolution patches with a tiling-and-stitching approach. The chosen 512×341 resolution thus represents a pragmatic balance between retaining sufficient visual information and achieving fast, CPU-friendly processing. This uniform reduction in size dramatically lowers memory usage and inference time, enabling the use of more advanced models or deployment on lower-capability hardware. As a result, the final model can process images at roughly one frame per second on a standard CPU, which enabled us to release a free, real-time demo via Hugging Face (Fernandez and Atef, 2025) running online on non-specialized hardware and publicly accessible without subscription.

We separated the test set at the start to avoid any leakage and built it using a random sampling of 236 images, spanning all the image providers. The remaining dataset was used as a training set, which was augmented before training and hyperparameter fine-tuning. Because the root cross-sections exhibit radial symmetry, we applied extensive rotation-based data augmentation. Each original image-mask pair was augmented with 24 rotated copies at regular 15° intervals (covering rotations from 0° to 345°). These rotations are biologically valid for cross-sections, which have no inherent “upright” orientation, effectively multiplying the training samples without introducing spurious features. Additionally, we applied horizontal and vertical flips, as well as small random translations (±10 pixels). This multi-faceted augmentation strategy expanded the training dataset to hundreds of thousands of images, providing an extensive set of examples to improve the model’s robustness.

### 2.6 SegFormer architecture and training procedure

We selected a vision transformer architecture and adapted it to our requirements by designing the output segmentation head to produce two output channels corresponding to the cortex-endodermis region (CE) and the lacunae/aerenchyma class (AR). We used the SegFormer 2D semantic segmentation framework (Xie *et al*., 2021), which combines a hierarchical Mix Transformer (MiT) encoder with a lightweight all-MLP decoder. SegFormer was originally introduced and benchmarked on natural-image segmentation datasets such as ADE20K, while its encoders are commonly initialised from ImageNet-1K pre-training before fine-tuning. Its design is well-suited to our task because it produces multi-scale feature maps that capture both global root organisation (cortex outline and tissue interfaces) and small-to-medium structures with high morphological variability (individual lacunae). In addition, SegFormer does not use explicit positional encodings, avoiding the need to interpolate positional embeddings when inference resolution differs from training. This is convenient in our workflow, where images are systematically resized for computational tractability and CPU-friendly deployment.

SegFormer is declined in multiple versions, using a family of backbones of increasing capacity (from B0 to B5). We evaluated variants B0 to B4 in preliminary experiments and selected SegFormer-B4 as an acceptable performance-speed compromise for final training. Models were trained in PyTorch on the Jean Zay supercomputing facility (GENCI, France) using NVIDIA H100 GPUs. Unless otherwise stated, training hyperparameters were: batch size 8, input size 512×512, AdamW optimiser, learning rate 1×10^−4^ with cosine decay, and weight decay 0.01. We used a combined loss function,

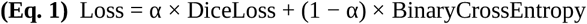

with α ∈ {1, 0.9, 0.7, 0.5} to assess robustness to loss weighting. Models were trained for up to 100 epochs with early stopping based on validation performance. Training logs were recorded using

TensorBoard, with reported metrics including standard metrics for segmentation tasks (Taha and Hanbury, 2015), such as IoU.

### 2.7 Evaluation of the pipeline

#### Automatic evaluation

Training logs were recorded using TensorBoard throughout all experiments. Model performance was first quantified by tracking both the evolution of global loss and Intersection over Union (IoU) for cortex– endodermis, CE, and lacunae, AR. After training, we assessed qualitative performance on a test set of images unseen by the model, drawn from all acquisition sites and experimental conditions, and summarized the results by reporting epoch-wise IoU on the training and validation set, and final Lacuna-to-Cortex Ratio R^2^ for each use-case (hydroponics, shelter, field) and across the whole population.

#### Independent expert review

We complemented these automatic metrics with an independent expert review to investigate the biggest errors and draw conclusions about the capability of the model to stick to the objective and to capture the variability of the target structures. First, we ranked all test images according to a disagreement score between manual and model masks (maximizing 1-IoU). From this ranking, we randomly sampled 23 images among those with the largest disagreements, corresponding to cases where at least one of the two segmentations (manual or model) was likely to be substantially incorrect. For each selected image, we identified the segmented aerenchyma components contributing to the discrepancy (connected pixel regions of lacuna that differed between masks) and presented them to the evaluators. Two expert rice anatomists not involved in the initial annotation independently inspected the raw images together with the manual and model segmentations. For every discrepant aerenchyma component, they assigned one of three labels: (i) error in the manual annotation, (ii) error in the model prediction, or (iii) undefined. The final counts are reported in the Table 2.

**Table 2.**
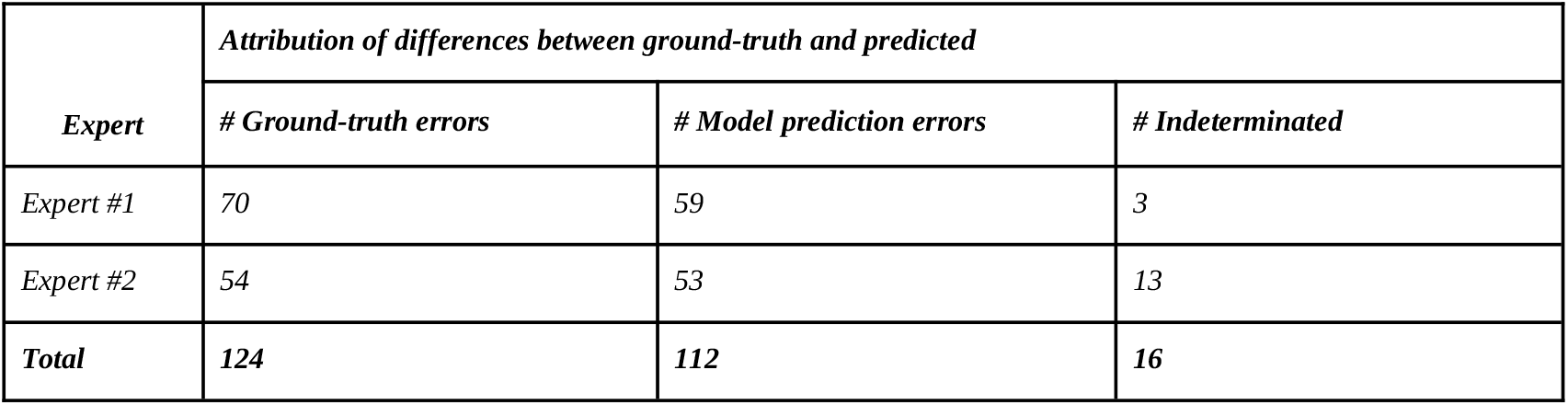
Results of the expert review of the model errors.

| Expert | Attribution of differences between ground-truth and predicted |  |  |
| --- | --- | --- | --- |
|  | # Ground-truth errors | # Model prediction errors | # Indetermined |
| Expert #1 | 70 | 59 | 3 |
| Expert #2 | 54 | 53 | 13 |
| **Total** | **124** | **112** | **16** |

### 2.8. Visibility, accessibility, and reproducibility

To ensure reproducibility and facilitate adoption by the community, all code used for preprocessing, training, inference, and visualisation is released under an open-source licence on GitHub, tagged v1.0.1, and secured with a Zenodo DOI (Atef H., 2025). The multi-environment multi-site dataset used for training can be shared in an anonymized format upon reasonable request. The automated processing pipeline is publicly available online as a HuggingFace space (Fernandez and Atef, 2025), together with documentation, a small dataset sampled from the test set images, and an online interactive demonstrator. The demonstrator allows non-specialists to use the demo dataset or to upload their custom batch of images stored in a .zip archive and obtain predictions in a limited time (∼2 seconds per image) without any local installation.

## RESULTS

### 3.1 Performance of SegFormer-B4 for aerenchyma and cortex segmentation

Across most hyperparameter configurations, the SegFormer-B4 model converged rapidly and consistently, with a high IoU reached on the validation set (Fig. 4, upper panel) after a few epochs. Across the four best configurations, IoU value on the validation set reached 0.90 after 3 epochs, 0.92 after 5 epochs, and its final best value after 10 epochs. Final mean validation IoU values ranged from 0.925 to 0.929 for lacunae and from 0.943 to 0.944 for cortical tissues (CE), with minimal variability across loss-weighting settings, indicating stable optimization and consistent performance over training experiments. In the training set curves (Fig. 4, bottom panel), the loss profiles (right plot) showed a smooth decline during the first 10 epochs and mostly plateaued thereafter, in conjunction with the trend change on the validation loss (upper-right plot) that indicates the start of training set overfitting and the loss of generalization capacity. Some variations were identified in the exact training curves trends, while the final loss obtained after training showed very small variations in final IoU across configurations. In the end, we selected the model with the best final Lacuna IoU value, while the small variations observed between the different models (worst=0.926, best=0.929) can be considered as having limited impact.

**Fig. 4.**
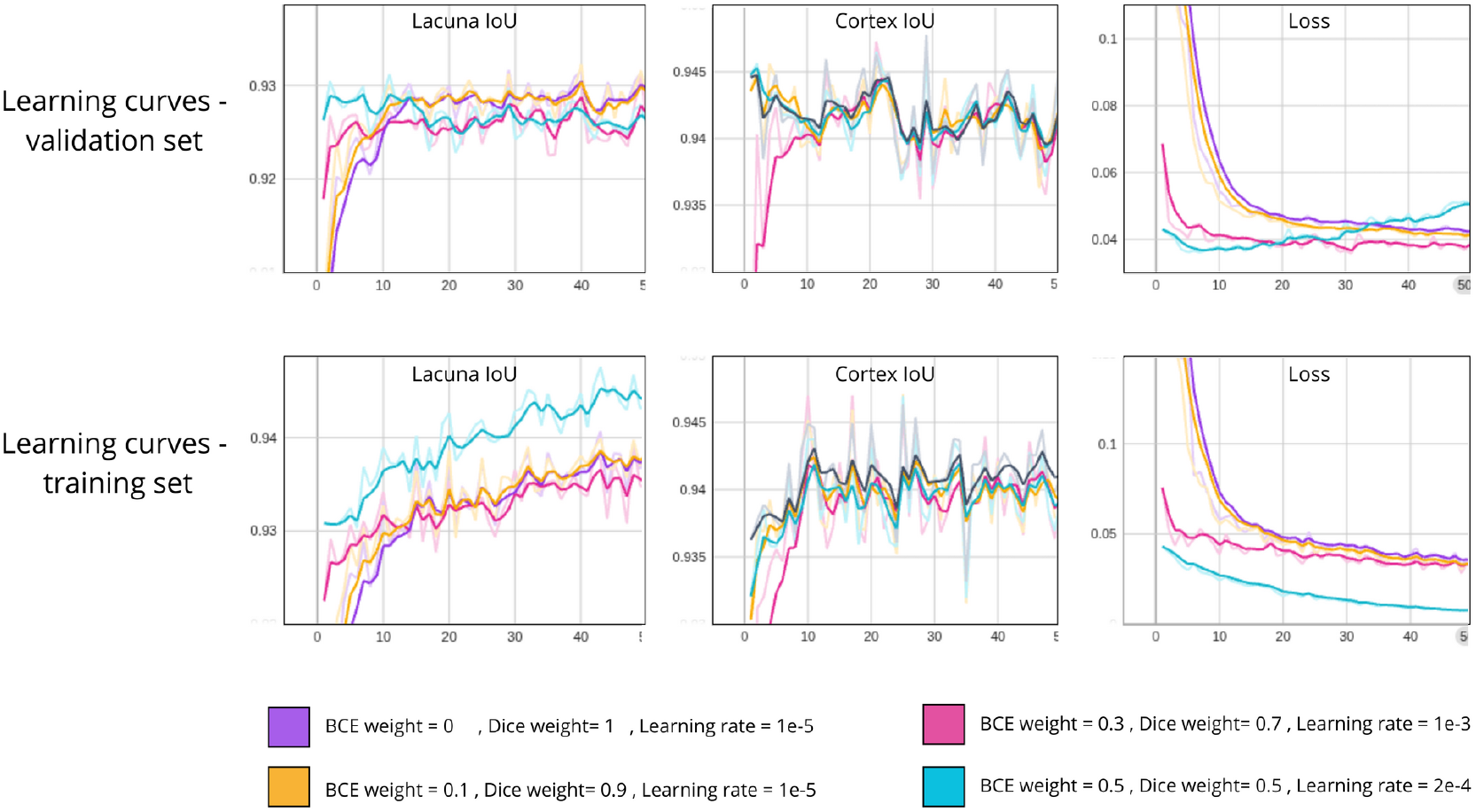
Training and validation performance (y-axis) versus training epochs (x-axis) across loss configurations for cortical and lacuna segmentation. Training and validation curves are shown for both cortex segmentation and aerenchyma lacunae segmentation, illustrating convergence behaviour under different loss formulations. **ALT-TEXT:** Polyline plots in the style of learning curves, showing training and validation performance as a function of training epochs for different loss configurations. Separate curves are displayed for cortex segmentation and aerenchyma lacunae segmentation, illustrating convergence behaviour and stability across loss formulations.

### 3.2 Performance of the full pipeline for estimating the Lacuna to Cortex Ratio

To evaluate the model’s ability to quantify aerenchyma and feed these traits to modeling experiments or breeding strategies, we analyzed the capacity of the pipeline to predict the target trait of interest (lacuna-to-cortex ratio, LCR), comparing the LCR value obtained with the predicted segmentations with the LCR value obtained from the ground-truth manual annotation (see Fig. 5). Because breeding captures trait variations across large populations, we investigated the capability of the model to generalize its training by predicting the variations of LCR on unseen data. To that end, we computed the coefficient of determination (R^2^) between predicted and ground-truth LCR values on the test set images. The coefficient computed on the full test set was very high, indicating a very good compliance with the expected objective (R^2^ = 0.98). We then separated the data by providers (six use-cases) and measured the R^2^ within each subset, obtaining consistently high values, which indicated high segmentation performance across use-cases.

**Fig. 5.**
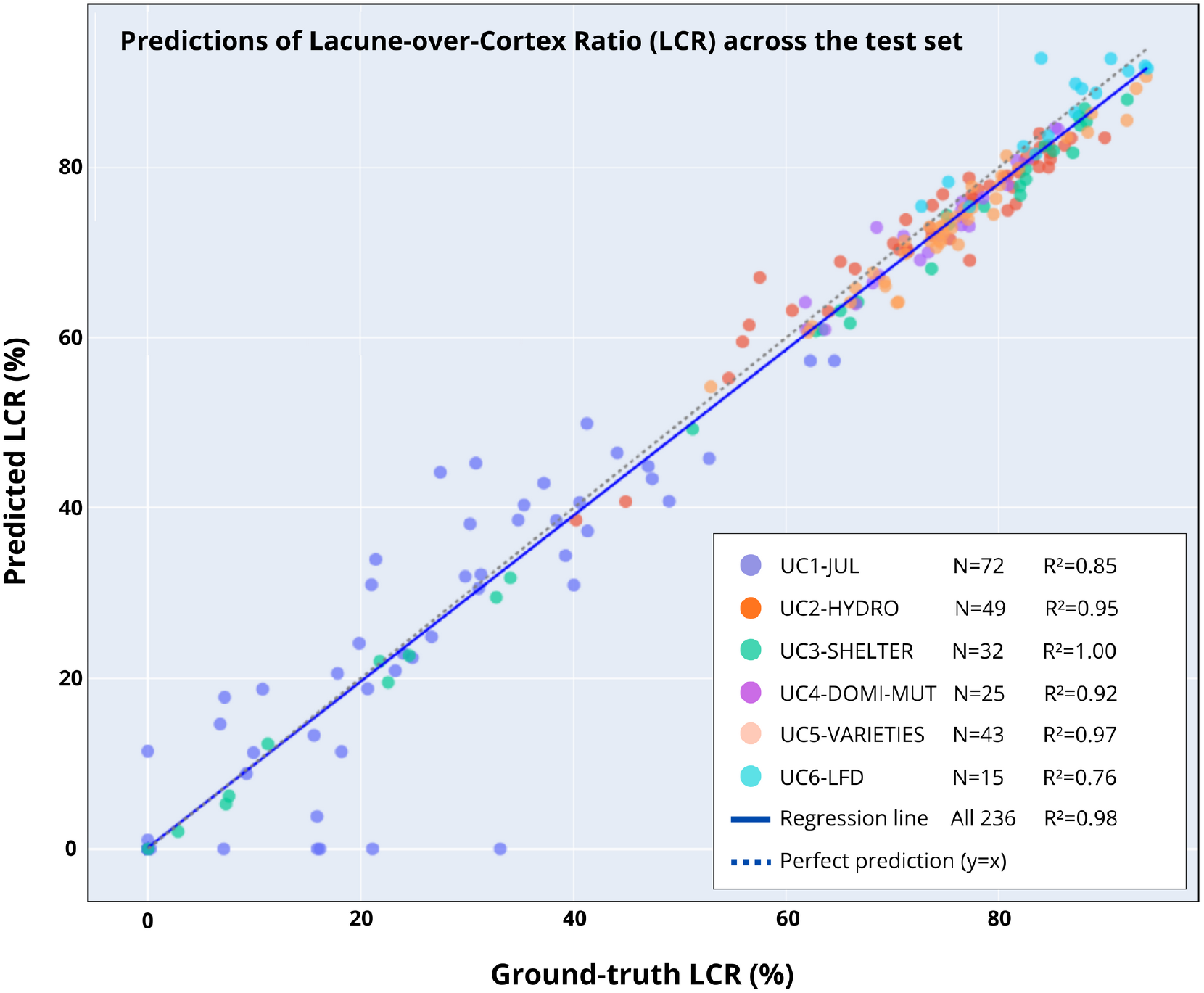
Agreement between predicted and ground-truth lacuna-to-cortex ratio. Each point represents one image (N = 236), coloured by acquisition use-case. The correspondence between predicted measurements (y-axis) and measurements from manual annotation (x-axis) indicates trait quantification performance across heterogeneous datasets. **ALT-TEXT:** Scatter plot comparing predicted and manually annotated lacuna-to-cortex ratios, with six different colours to identify the corresponding use-case.

Over the six use-cases, two exhibited an R^2^ lower than 0.93. Use-case 6 (UC6-LFD) had the lowest R^2^ value (0.76). However, this result has limited significance since the subset contains only 15 images, with a high LCR mean (0.8) and a small standard deviation around the mean (0.1). In this context, the range of variations is very small relative to the mean value, making small errors appear disproportionately impactful on the R^2^ values.

The second “difficult” use-case (UC1-JUL) had a larger number of images and an LCR range covering most of the overall variability of LCR across the population, while exhibiting a relatively low R^2^ at 0.85, which can be considered as an indicator of the real difficulty of the pipeline to stick to the objectives in this dataset. Visual inspection by independent experts showed that this use case included a high number of young roots, and therefore more early-stage aerenchyma, which are harder to annotate, and that it was affected by more misinterpretations of the annotation guide. We described this aspect below, presenting example images and metrics in the results section 3.4, “Independent expert review”.

### 3.3 Qualitative assessment of segmentation

Visual inspection confirmed that the model reliably captured the shape and position of lacunae across a broad range of anatomical configurations. In clear and well-contrasted images, predicted lacuna masks closely matched ground-truth delineations, capturing even small cavities. In more challenging images, especially those with blurred cortical boundaries or irregular staining, the model tended to slightly overestimate lacunae shape, but still preserved their general geometry and did not introduce anatomically implausible structures (*e*.*g*., lacuna outside the cortex, or scattered parts of lacunae). Unfrequently, segmented images displayed extremely bad predictions. Most often, these unsuccessful cases combined several challenges described earlier (blurry images, low cell wall contrast), with additional difficulties, such as ambiguous lacunae corresponding to young and undifferentiated aerenchyma, which induced annotation issues (see Fig. 6A).

**Fig. 6.**
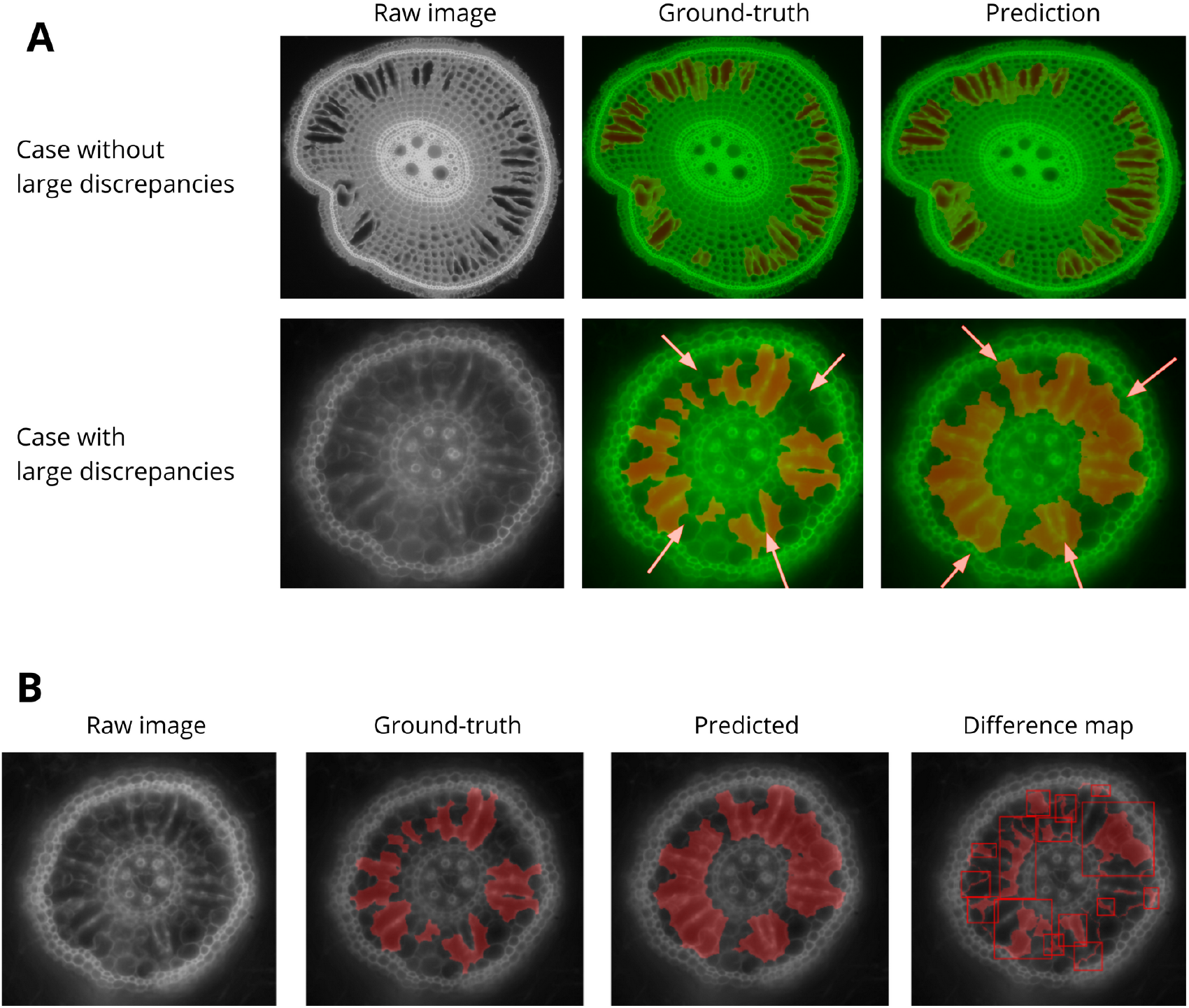
A: qualitative examples of automatic segmentation performance on representative root cross-sections. Left: raw data. Middle and right: overlay of raw data (green channel) and segmentation output (red channel). The upper panel shows a high-quality automatic segmentation with accurate identification of aerenchyma lacunae. The lower panel illustrates a failure case, with large discrepancies between the ground truth and the predicted lacunae. B: Expert review of ground-truth versus model errors assisted by an algorithm to highlight the discrepancies. **ALT-TEXT:** Figure illustrating qualitative segmentation performance and expert evaluation. Panel A shows representative root cross-sections with raw images and overlays of raw data and segmentation output, highlighting an accurate segmentation example and a failure case with discrepancies relative to the ground truth. Panel B presents an expert-assisted comparison of ground-truth annotations and model predictions, with algorithmically highlighted differences to support error assessment.

### 3.4 Independent expert review

Visual inspections showed that some of the discrepancies between the ground truth and the model predictions could probably be attributable to annotation errors (Fig. 6B). We further investigated this intuition by constituting an additional panel of two expert biologists to review a random set of images showing large discrepancies between ground-truth and predictions (23 images selected among those with the lowest IoU prediction values). Interestingly, a majority of these images were associated with UC1-JUL experimental set, which had the lowest R^2^ score.

#### Quantitative analysis

We provided the experts with the raw images, ground-truth masks, model predictions, and difference maps (ground-truth versus predictions). For each mismatch (in the general case, there are multiple mismatches per image), they attribute it to the model, the annotator, or undetermined. These results are presented in Table 2. Across all reviewed images, experts identified 124 ground-truth annotation errors versus 112 model prediction errors. This indicates a higher number of errors made by annotators (+11 %) than by the model. In addition, 16 discrepancies (6 % of the differences) could not be attributed, suggesting human perceptual limits or remaining limitations in the problem definition.

#### Qualitative analysis

We then asked the experts to provide a qualitative assessment of the dataset. Importantly, experts noted that manual annotation errors typically corresponded to large, misplaced regions with a substantial impact on the computed lacuna percentage, whereas model errors were generally restricted to thin boundary shifts or local contour irregularities. These results support the view that the network at least matches, and sometimes exceeds annotators by avoiding major inconsistencies that arise probably from human subjectivity, objective misunderstanding, or fatigue.

### 3.5 Application to rice root phenotyping across environments, genotypes, and growth stages

#### Deployment in biology

The automated segmentation pipeline provide accurate aerenchyma segmentation while reducing per-image processing time from approximately 3 minutes per slice (semi-automatic workflow) to almost two seconds on CPU, and even less on GPU. This gain enables routine analysis of large anatomical datasets, removes operator-dependent variability, and allows breeding teams to quantify aerenchyma at scales previously inaccessible. The model and codebase have been adopted by collaborating biologists and integrated into ongoing phenotyping campaigns, starting from the 6 use-cases covered in this study

#### Overview of usage across use cases

The pipeline is currently used across six complementary studies that span gene discovery, stress physiology, and breeding-oriented screening. UC1 screens rice mutants to identify genes associated with contrasting trajectories of cortical aerenchyma development. UC2 (Colombia) evaluates a diversity panel of cultivated rice varieties under hydroponics designed to mimic flooded paddy conditions, to characterize anatomical responses to low-oxygen environments. UC3 (Colombia) assesses varietal responses under controlled irrigation regimes in the field (shelter), quantifying aerenchyma plasticity under contrasting water supply. UC4 extends UC1 to soil-like growth conditions to test whether mutant phenotypes observed in controlled settings persist in more realistic substrates. Finally, UC5 (France) and UC6 (Colombia) screen diverse germplasm panels to map aerenchyma profiles and provide trait distributions that can inform upcoming breeding efforts.

#### Focus on an ongoing work in use-cases (UC2 and UC3)

Aerenchyma formation in roots is a key anatomical trait associated with how plants respond to varying levels of soil oxygen availability (Pedersen *et al*., 2021). Therefore, quantifying the characterization of root anatomical traits related to aerenchyma is important for understanding the functional plasticity of the root system in contrasting environments. Root anatomical traits were quantified 1 cm proximal to the root plateau using the phenotyping pipeline across multiple experimental contexts (Fig. 7). The lacuna-to-cortex ratio (%) differed significantly between flooded and non-flooded conditions across genotypes, with significant genotype and genotype × environment effects. Cortex surface area differed between treatments only under hydroponic conditions, whereas stele surface area differed only under soil conditions. Genotypic variation was greater for cortex and stele surface area than for the lacuna-to-cortex ratio. For both cortex and stele surface area, genotype and genotype × treatment interactions were significant. (Fig. 7A).

**Fig. 7.**
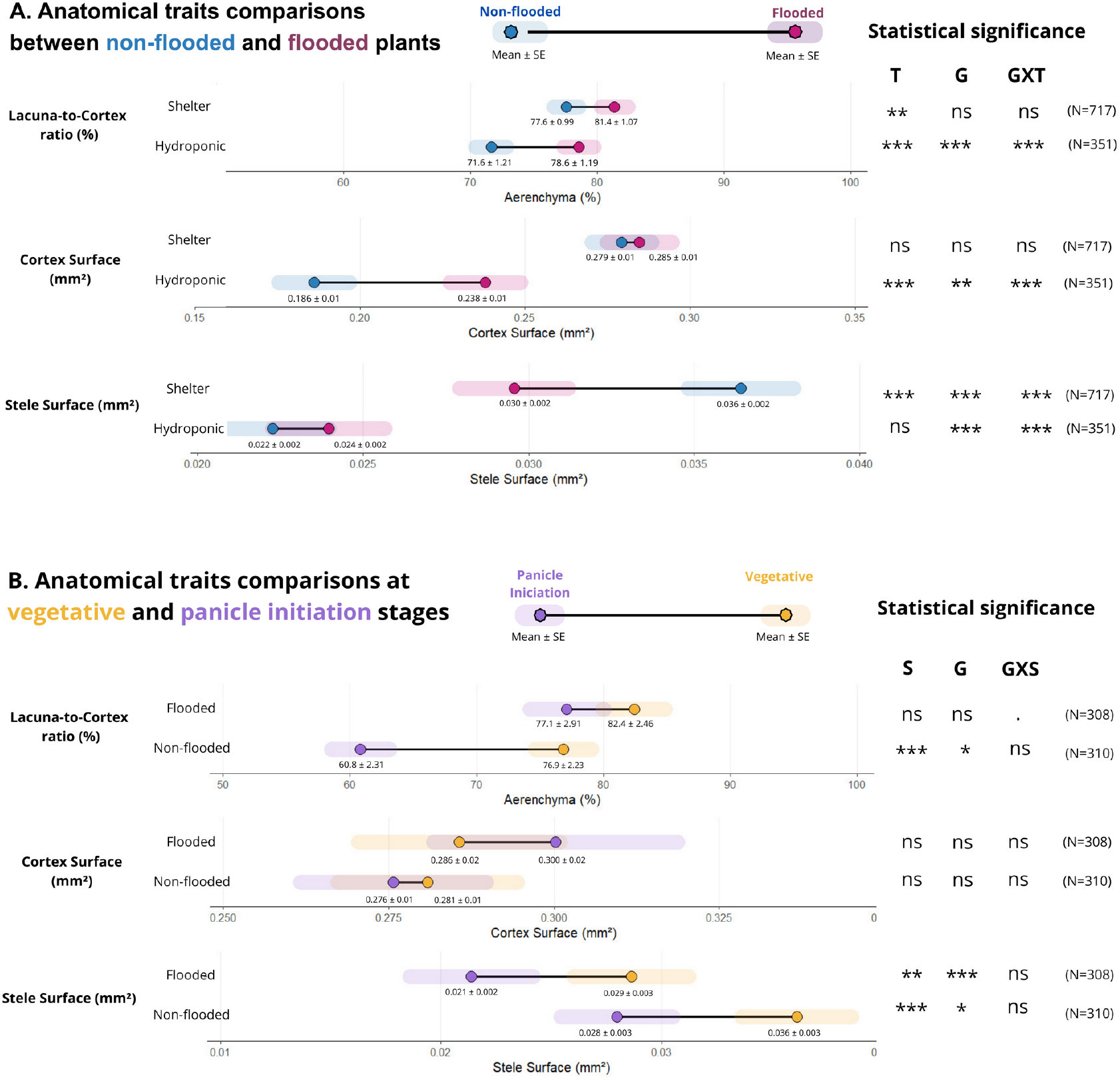
Root anatomical traits variations across environmental and developmental factors in experimental systems with different levels of oxygen availability. Points represent estimated marginal means (EMMs), with horizontal bars indicating ± standard errors, and solid lines represent pairwise contrasts. Statistical significance was assessed using two-sided mixed-model ANOVA, and main effects and interactions are reported on the right side of each panel dusing standard significance codes: ns, not significant; 0.05 ≤ P 0.10 (.); P < 0.05 (*); P < 0.01 (**) and P < 0.001 (***). Tested effects include genotype (G), treatment (T), developmental stage (S), and their interactions (GxT, GxS). Each panel reported the sample size (N) corresponding to the number of root cross-section images analyzed per experimental group, obtained from independent plants (biological replicates). A: trait comparisons between non-flooded and flooded conditions under field (Shelter) and hydroponic experiment conditions. B: trait comparisons between vegetative and panicle initiation developmental stages under contrasting water regimes in field conditions. **ALT-TEXT:** Multi-panel figure showing variation in rice root anatomical traits across environmental conditions and developmental stages. Points represent estimated marginal means with associated uncertainty, and pairwise contrasts illustrate treatment and stage effects. Panels compare flooded versus non-flooded conditions and vegetative versus reproductive stages across experimental systems, highlighting the influence of genotype, environment, and development on root anatomy.

To assess the ability of the tool to capture developmental effects on root cortical traits, anatomical variables were compared between the vegetative stage and panicle initiation under shelter conditions (Fig. 7B). Under non-flooded conditions, the proportion of aerenchyma in the cortex was significantly higher at panicle initiation than at the vegetative stage (P < 0.001), whereas no developmental differences were detected under flooded conditions. Cortex surface area did not differ significantly between developmental stages under either water regime (P > 0.05). In contrast, stele surface area varied significantly with development under both flooded (P < 0.01) and non-flooded (P < 0.001) conditions, with consistently larger stele surface area at the vegetative stage. Significant variation in cortex and stele surface area was observed within each treatment and developmental stage, driven by significant genotypic effects.

To evaluate the genotypic variation structure by ecology within a single environment, the distribution of lacune-to-cortex ratio (%) was examined across rice ecological groups under aerobic field conditions (Fig. 8). There were no significant differences in overall mean aerenchyma in cortex (%) values among ecologies (Type III ANOVA, P > 0.05), without significant contrast between ecological groups after Tukey asjustment (P > 0.05 for all comparisions). Despite this, Lowland, Lowland-upland, and Upland genotypes exhibited partially overlapping distribution, with all ecological groups spanning a wide range (≈54-96%). This reflects substantial variability within each ecology under uniform conditions.

**Fig. 8.**
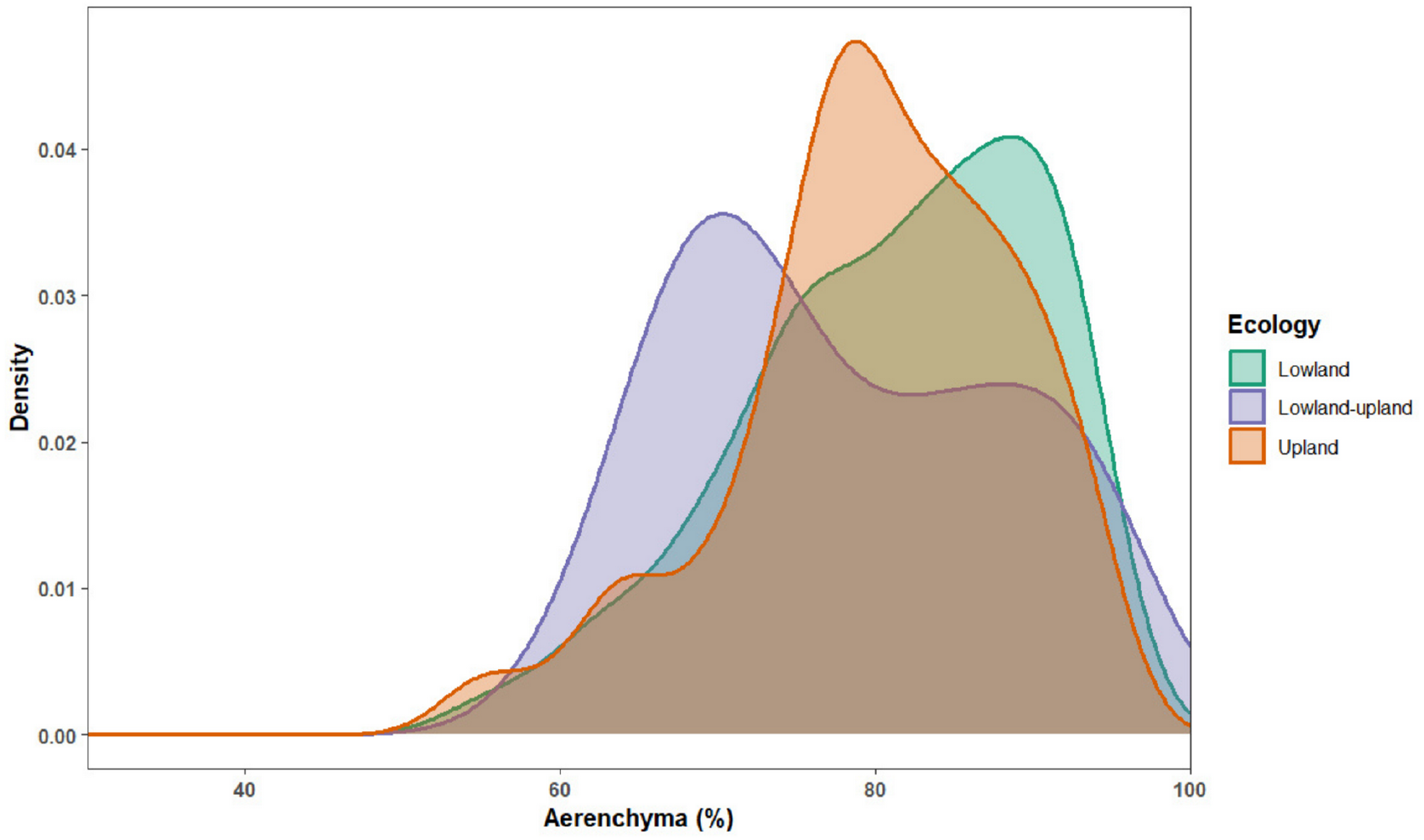
Density distributions of lacuna-to-cortex ratio (%) across ecological groups of rice under aerobic field conditions. Curves density show the frequency distribution of the aerenchyma values measured in 40 genotypes grouped by ecology (Lowland, Lowland-upland, Upland). Differences in curve position, width, and overlap reflect contrasting ecological variation in root anatomical traits in response to non-flooded growth conditions. Measurements were obtained from independent plants (biological replicates) grown under non-flooded conditions. Sample size (N=600) corresponds to the total number of root cross-section images analyzed. **ALT-TEXT:** Density plots showing the distribution of lacuna-to-cortex ratio values across rice ecological groups under aerobic field conditions. Separate curves represent lowland, lowland–upland, and upland groups, illustrating differences in distribution shape and overlap that reflect contrasting anatomical variation among ecologies.

Across ecological groups, lacune-to-cortex (%) exhibited a consistent ordering of mean values, with higher averages in Lowland genotypes (mean ± std-deviation: 81.1 ± 9.34, N = 255), intermediate values in Upland genotypes (79.9 ± 10.3, N = 300), and lower values in Lowland-upland genotypes (77.9 ± 8.98, N = 45). However, density distributions (Fig. 8) revealed that Upland genotypes were characterized by a strong concentration of observations around intermediate values (≈70%), whereas Lowland genotypes displayed a distribution encompassing both moderate and high values more centered on higher values.

In this way, the tool allows for demonstrating the variability across genotypes for aerenchyma formation in the shelter experiment and differences between ecologies.

## DISCUSSION

### A high-throughput, open, and transferable anatomical phenotyping tool

This study demonstrates that a recent vision transformer architecture enables robust, high-throughput segmentation of rice root anatomy under heterogeneous experimental and imaging conditions. Automated phenotyping of root anatomical traits is increasingly recognized as a major bottleneck in root biology and breeding, and existing tools often rely on semi-automatic workflows, site-specific tuning, or restrictive acquisition conditions (Burton *et al*., 2012; Chopin *et al*., 2015; Lartaud *et al*., 2015; Atkinson *et al*., 2019; Wang *et al*., 2020). By contrast, our SegFormer-based pipeline was trained on a multi-environment dataset encompassing field-derived and controlled images, and did not require any site-or condition-specific calibration to maintain high performance. However, we identify an inherent limitation to our evaluation process: while the test set images are “unseen data” for the model (they were not used for training), these images were acquired along with the images of the train set, during the same experiments. An additional level of validation would be to run a new validation step with data from new experiments, and ideally with images from other providers.

The combination of a lightweight decoder and a resolution-agnostic transformer encoder allowed efficient inference while preserving anatomical detail, making the method compatible with large-scale deployment and CPU-accessible usage. This design choice supports practical phenotyping workflows, where thousands of images from multiple sources must be processed rapidly and reproducibly. The open release of code, trained models, annotation guidelines, and an interactive online demonstration further ensures transparency, reproducibility, and reusability, aligning with current recommendations for AI-driven plant phenotyping tools (Ubbens and Stavness, 2017; Rädsch *et al*., 2023). In practical terms, the pipeline’s scalability has already been demonstrated by its deployment in ongoing phenotyping campaigns, processing thousands of images from multiple field sites, with minimal human intervention.

### Expert-level precision and reduced subjectivity relative to manual annotation

Beyond throughput, our results highlight a critical advantage of deep learning for anatomical phenotyping: increased consistency relative to human annotations. While expert manual segmentation remains the reference standard, it is inherently limited by subjectivity, fatigue, and ambiguity in class boundaries, particularly for fine or developing aerenchyma lacunae. Annotation variability is well documented in bioimage analysis and can substantially hinder both model training and biological interpretation (Rädsch *et al*., 2023; Taha and Hanbury, 2015).

In our study, an independent expert audit showed that model predictions matched or exceeded the reliability of individual human annotators. Notably, the model handled challenging cases with greater consistency than individual human annotators. Model errors were primarily limited to subtle boundary shifts, whereas manual annotations occasionally exhibited larger inconsistencies, especially in datasets dominated by young roots with small or not-well differentiated lacunae. We interpret this behaviour as a consequence of training on pooled expert knowledge across datasets, which enforces anatomical coherence and reduces individual bias. In this sense, as shown in the biomedical imaging community using multi-expert learning frameworks such as GroupLearning (Cao *et al*., 2024), models trained on annotations from multiple experts can mitigate individual rater bias and yield more stable and reproducible predictions for tasks affected by perceptual ambiguity.

However, our results suggest that further improvement in lacuna segmentation accuracy could be achieved by: i) improving annotator’s guidance on young aerenchyma, for example, by complementing the annotation guide, having a round-table by sharing the first 5% images annotated), ii) increasing the number of annotated images, as current training set relies on data augmentation (random rotations, translations and mirror operations from a handful of images, roughly 1700), and iii) building on these improvements, by exploring additional training configurations, alternative architectures (heavier than SegFormer-B4, at the expense of deployability) or redesign the training workflow using annotation-efficient approaches such as active learning.

#### Discrimination of root cortical traits across genotypes, environments, and developmental stages

Automated segmentation translated into biologically meaningful trait quantification. The AI-based framework consistently discriminated against genotypic effects, environmental responses, and developmental shifts in cortex and stele surfaces as well as lacuna-to-cortex ratios across hydroponic and soil-based systems. These results demonstrate that automated image analysis preserved known anatomical responses while revealing fine-scale variation across contexts.

Increased aerenchyma formation under flooded or oxygen-deficient conditions was detected in both hydroponic and shelter experiments, consistent with established physiological mechanisms driven primarily by oxygen availability rather than soil factors per se (Justin and Armstrong, 1991; Colmer, 2003; Pedersen *et al*., 2021). Differences observed between hydroponic and soil-grown roots further suggest an influence of mechanical impedance on cortical and stele development, in line with previous reports on root anatomical plasticity (Lynch *et al*., 2021; Jones *et al*., 2025b; Qu *et al*., 2008). The tool also captured developmental stage effects, including shifts in aerenchyma proportion and stele size between vegetative and reproductive stages, reinforcing its sensitivity to growth-related anatomical dynamics (Chen *et al*., 2021; Jones *et al*., 2025a).

The ability to quantify large genotypic variance in cortical traits (Figure 8) highlights the relevance of AI-enabled phenotyping for breeding and diversity studies. Root cortical aerenchyma is increasingly recognized as a key trait mediating adaptation to contrasting water regimes and contributing to methane emission dynamics in rice systems (Suralta and Yamauchi, 2008; Jiménez and Pedersen, 2023; Caetano *et al*., 2025). Yet, despite its importance, explicit selection on cortical aerenchyma remains unfrequent in breeding programs (Jones *et al*., 2025b). By enabling reproducible, multi-environment anatomical phenotyping at scale, our pipeline provides a foundation for integrating rice root anatomy into physiological modelling, genetic mapping, and genomic selection frameworks.

Overall, this work demonstrates that transformer-based image analysis can deliver scalable, precise, and biologically informative root anatomical phenotyping under field conditions. By addressing annotation variability and enabling cross-environment trait quantification, the proposed approach supports more reproducible comparisons and accelerates efforts to understand and exploit root anatomical plasticity for crop adaptation and climate-smart rice breeding.

## ACKNOWLEDGEMENTS

This work was carried out using HPC resources from GENCI at Saclay, France, under allocation AD011012561R4. We acknowledge the imaging facility MRI, a member of the national infrastructure France-BioImaging, supported by the French National Research Agency (ANR-24-INBS-0005 FBI BIOGEN). The authors thank all colleagues and technical staff who contributed to plant cultivation, sample preparation, microscope acquisitions, and experimental support. We also acknowledge valuable discussions and logistical assistance that helped facilitate this study. Special thanks to #DigitAg “Digital Agriculture Convergence Lab” (www.hdigitag.fr) for its supportive community.

## AUTHOR CONTRIBUTIONS

HA, BC, CP, MCR, and RF: conceptualization; HA, LFD, PALM, SNS, JB, BC, CP, MCR, and RF: methodology; HA and RF : software; HA, LFD, PALM, MCR, and RF: formal analysis; MCR and RF: validation; HA, LFD, PALM, SNS, JB, BC, CP, MCR, and RF: investigation; CP, MCR, and RF: resources; HA, LFD, PALM, SNS, and JB: data curation; HA, LFD, BC, CP, MCR, and RF: writing - original draft; HA, LFD, BC, CP, MCR, and RF: writing - review & editing; HA, LFD, MCR, and RF: visualization; BC, CP, MCR, and RF: supervision; BC, CP, MCR, and RF: project administration; BC, CP, MCR, and RF: funding acquisition

## DATA AVAILABILITY

The datasets generated and analyzed in this study include 1760 images of raw mmicroscopy images and the corresponding annotations. Due to the large volume of these files and current storage and transfer constraints, the data are available from the corresponding author upon reasonable request. To ensure reproducibility and facilitate adoption by the community, all code used for preprocessing, training, inference, and visualisation is released under an open-source licence on GitHub, tagged v1.0.1, and secured with a Zenodo DOI (Atef H., 2025). The multi-environment multi-site dataset used for training can be shared in an anonymized format upon reasonable request. The automated processing pipeline is publicly available online as a HuggingFace space (Fernandez R., 2025), together with documentation, a small dataset sampled from the test set images, and an interactive inference demonstrator. The online demonstrator allows non-specialists to upload a batch of images stored in a .zip archive and obtain predictions in a limited time (∼2 seconds per image) without any local installation.

